# JACKS: joint analysis of CRISPR/Cas9 knock-out screens

**DOI:** 10.1101/285114

**Authors:** Felicity Allen, Fiona Behan, Francesco Iorio, Kosuke Yusa, Mathew Garnett, Leopold Parts

## Abstract

Genome-wide CRISPR/Cas9 knockout screens are revolutionizing mammalian functional genomics. Their range of applications remains limited by signal variability from different guide RNAs targeting the same gene, which confounds analysis, and dictates large experiment sizes. To address this problem, we report JACKS, a Bayesian method that jointly analyses screens performed with the same guide RNA library. Modeling the variable guide efficacies greatly improves hit identification, and allows a 2.5-fold reduction in required cell numbers without sacrificing performance compared to current analysis standards.

---

CRISPR/Cas9 knockout screens can assess the influence of every gene’s knockout on any selectable cellular trait in a single assay^1, 2^. The guide RNA (gRNA) libraries used in these experiments typically contain several gRNAs per gene, each recruiting the Cas9 protein to inflict a loss-of-function mutation. Genes required for the selected trait are mapped by introducing the gRNA library into cells, applying selection, sequencing the gRNA locus, and processing the data using methods such as MAGeCK^3^ or BAGEL^4^.

A central source of confounding in analysis of screen outputs is conflicting evidence from alternative gRNAs targeting the same gene, caused by different gRNA efficacies^5^. This variability has been linked to a range of technical and biological factors^6–11^, and while several gRNA efficacy estimation algorithms have been proposed^2, 7, 12–14^, their predictive ability remains limited ^15, 16^. As a result, screens still use five or more gRNAs per gene, and at least three replicates are recommended^17^, rendering the required scale a bottleneck for systematic assessment of gene function and genetic interactions, particularly in short term primary cultures.

To overcome this issue, we present JACKS (Joint Analysis of CRISPR/Cas9 Knockout Screens), a Bayesian method that models gRNA efficacies in multiple screens that use the same gRNA library. Briefly, JACKS models log_2_-fold changes of gRNA read counts between treatment and control conditions as a product of treatment-dependent gene essentiality and treatment-independent gRNA efficacy (Figure 1A). We obtain approximate posterior probability distributions for these components, whilst accounting for experimental noise (Methods). The inferred gRNA efficacies match intuition (e.g. weaker effects due to gRNAs 1 and 6 in Figure 1A), are reproducible (Methods, Figure S1), and concordant with both CERES and Doench-Root efficacy scores^12, 18^ (Figure 1B, S2), supporting their use for improved screen analysis. The inferred gene essentialities provide a measure of the gene knockout’s log_2_-fold change in frequency between control and treatment conditions, corrected for noise and gRNA efficacy. For example, as expected, knocking out the KRAS gene was inferred to have a greater impact on growth in cell lines known to harbour KRAS driver mutations in the Aguirre dataset^19^ (Figure 1A). The variance of these estimates can be used to calculate a p-value of the effect.

**Figure 1:**
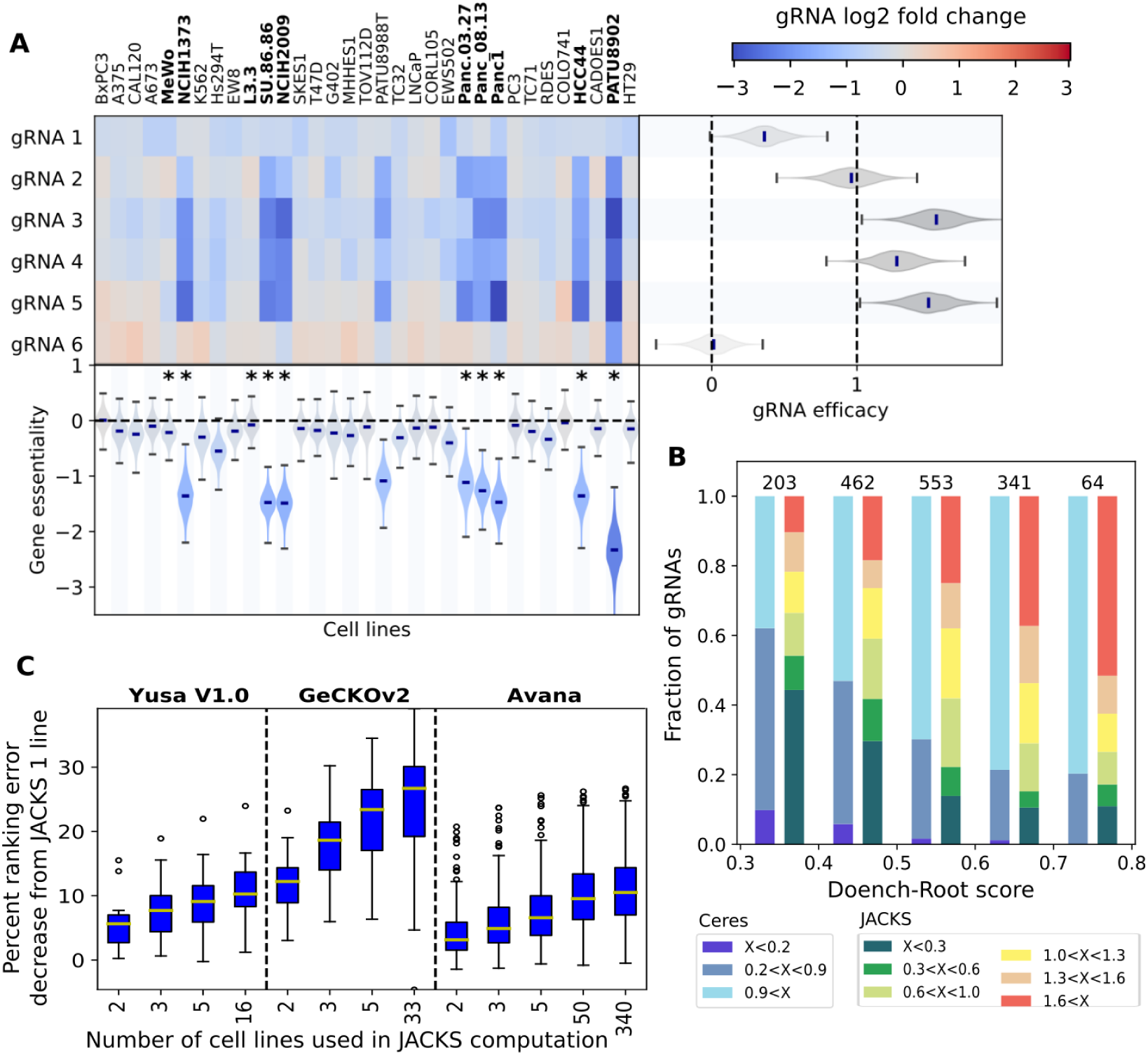
**A.** JACKS inferred decomposition of median-normalized log_2_ fold-change (heatmap) for six gRNAs targeting the KRAS gene (y-axis, GeCKOv2 library) in cancer cell lines from Aguirre *et al.*^19^ (x-axis). The inferred gRNA efficacies and gene essentialities (with uncertainty) are displayed to the right and below the heatmap, respectively. Lines with KRAS driver mutations are highlighted in bold and indicated with an asterisk. **B.** Fraction of gRNAs (y-axis) targeting Hart essential genes^20^ in each range of Doench-Root score^12^ (x-axis) for specified ranges of CERES and JACKS inferred gRNA efficacy scores (”x”, colors). Number of gRNAs in each column is marked above the bar. **C.** Percent ranking error (fraction of area above the ROC curve below 0.2 false positive rate, Methods) decrease (y-axis; median, quartiles, and 95th deciles marked in box plot) for increasing number of experiments in JACKS model (x-axis) for three different libraries.

To examine JACKS’s ability to identify screen hits, we used it to rank genes by their inferred essentiality, and evaluated how well this discriminates known essential from non-essential genes (defined by Hart *et al*^20^). We considered data from pooled knock-out screens performed with the Yusa v1.0 ^21, 22^, GeCKOv2^10, 19^, and Avana^23^ gRNA libraries, and measured performance using the 0.2 partial area under the curve (”Ranking accuracy”, Figure S3, Methods) and above the curve (”Ranking error”) metrics; equivalent results were obtained using alternative thresholds and criteria (Figure S4-S7, Table S1). Increasing the number of cell lines processed by JACKS from a single line to all available lines in each set reduced the median ranking error by 10%, 26% and 11% respectively (Figure 1C), with the first five additional lines providing the majority of the gains. While improvements were largest for the GeCKOv2 library, likely due to its lower starting performance and more variable gRNA efficacy^12, 23^, all three datasets benefited from joint screen analysis with JACKS.

We next compared the performance of JACKS to the most commonly used single-screen analysis methods; averaging the log_2_-fold-changes of all gRNAs targeting the gene (”MeanFC”), MAGeCK^3^ and BAGEL^4^. JACKS improved accuracy for 97%, 99%, and 91% of all cell lines tested, respectively, with a 12%, 21%, and 9% lower error on average (Figure 2A). To put these improvements in perspective, if there are 1,000 true hits to be recovered from roughly 19,000 genes tested, then in order to correctly identify the first 750 of them (0.75 recall), the other methods are expected to yield 684, 666, and 432 false positives on average, compared to JACKS’s 333 (Table S1). When applied to data from each cell line separately, the results for JACKS were equivalent to the alternatives (Figure S8). This demonstrates that although JACKS was designed to efficiently integrate information across experiments, there is no downside to using it on a single screen.

**Figure 2:**
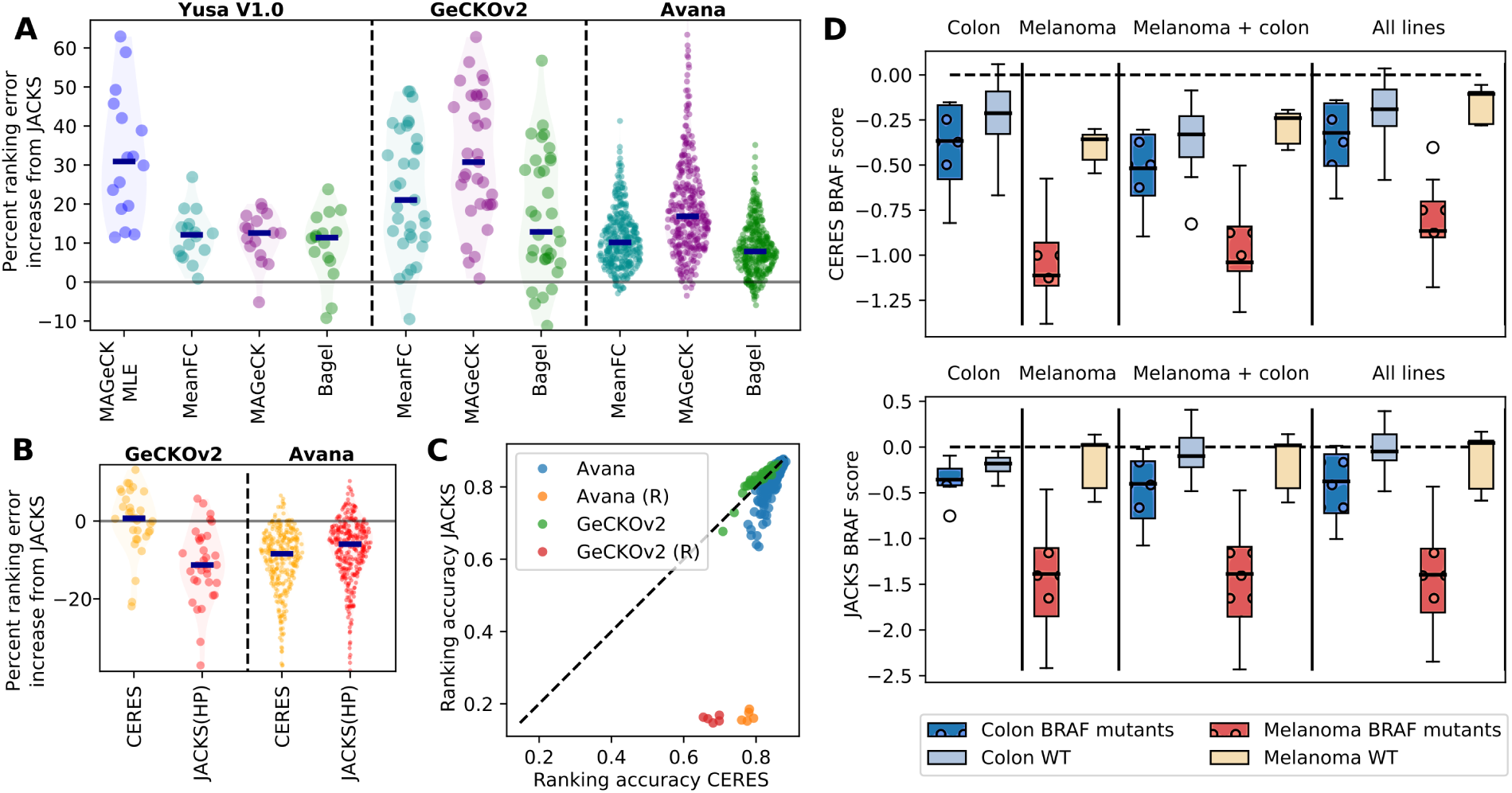
**A.** JACKS outperforms existing alternatives at distinguishing essential genes. Percent ranking error increase (y-axis) compared to JACKS for four alternative analysis methods (x-axis) on three different libraries (panels). Every marker represents one cell line; median increase is marked with a dark blue line, and estimated distributions are shaded in. **B.** Methods that assume similar gene essentialities across cell lines perform favourably compared to JACKS. Percent ranking error increase compared to JACKS (y-axis) for CERES (yellow) and JACKS with a hierarchical prior (HP) (red) for GeCKOv2 and Avana libraries. Markers and shading as in A. **C.** CERES identifies essential genes from random data. Ranking accuracy of CERES (x-axis) compared to JACKS (y-axis) on Avana (blue) and GeCKOv2 (green) libraries. Each marker corresponds to one cell line, with five randomized experiments (yellow and red markers) included for comparison. Dashed line, *y* = *x*. **D.** CERES’s preference for a common gene response across cell lines results in more similar scores for differentially essential genes, whereas JACKS maintains differential signal between cell lines. CERES (top) and JACKS gene essentiality scores for the BRAF gene in melanoma and colon cancer cell lines (colours) when processed with selections of cell lines (panels) from the Avana dataset, grouped by BRAF mutation status (shading and patterns).

Two existing methods jointly model outputs from multiple screens, MAGeCK-MLE^24^ and CERES^23^. We could run MAGeCK-MLE on only the smallest, Yusa v1.0 dataset due to large CPU and memory requirements (81 CPU days on 4-23 cores, 75 Gb of RAM for 16 screens), and observed it to offer no improvement over standard MAGeCK (Figure 2A). CERES performed equivalently to JACKS on GeCKOv2 data, and was more accurate on the Avana dataset (−0.7% and 8.4% lower median ranking error, Figure 2B).

The CERES model assumes some shared gene essentiality signal across experiments. To evaluate the impact of this model assumption, we introduced five additional screens into both the GeCKOv2 and the Avana datasets, each containing shuffled gRNA responses from a randomly selected cell line. CERES was able to identify essential genes from randomized data with high accuracy, while JACKS achieved the expected near-random performance (Figure 2C). We supplemented JACKS with an option to make a similar assumption of shared gene essentiality (JACKS(HP), Methods), and confirmed that this change resulted in comparable error to CERES on both GeCKOv2 and Avana data (Figure 2C), and correspondingly improved ability to extract signal from the lines with shuffled data (Figure S9).

Sharing gene effects across screens is beneficial for finding universal hits, but could mask true context-specific signal. To test this possibility, we examined BRAF essentiality in melanoma, where BRAF mutations are predictive of sensitivity to BRAF inhibitors^25^, and in colon cancer, where BRAF mutations are less prevalent, and predict only a weak response to BRAF inhibitors^26^. Accordingly, JACKS’s estimated BRAF essentiality in the Avana dataset is large in BRAF mutant melanoma lines, weak in BRAF-mutant colon cancer lines and negligible in most other lines (median −1.39 vs −0.35 vs 0.03, Figure 2D), regardless of the dataset used in estimation. CERES’s preference for a common gene response alters its estimates depending which other lines are selected for co-processing. BRAF essentiality score is lower in mutant melanoma lines when processed with all lines compared to when processed with melanoma lines alone (median −0.86 vs −1.11). Conversely, its essentiality is estimated to be larger in other lines when processed together with mutant melanoma lines instead of all data (median −0.36 vs −0.11, −0.52 vs −0.32 and −0.33 vs - 0.19 for non-mutant melanoma lines, non-mutant colon lines, and mutant colon lines, respectively, Figure 2D). Including melanoma lines with colon cancer lines in JACKS estimation increases the separation of BRAF mutants and non-mutants in the colon cancer lines (AUC 0.72 vs 0.78 in colon only, and colon+melanoma lines), suggesting that BRAF mutation status is still predictive in colon cancer, if only of a much weaker response.

Finally, we tested if improved analysis methods can be used to reduce experiment size without compromising findings. First, we considered the number of replicate screens. We performed twelve replicate experiments with the Yusa v1.0 library on the HT29 cell line (Methods), and combined these with the above described Yusa v1.0 dataset, which contains 2-3 replicates each for 15 additional cell lines. Identifying essential genes from three replicates of HT29 using JACKS, when co-processing with three replicates from each of the other 15 lines outperformed processing of the 12 HT29 replicates in isolation. Similarly, applying JACKS to just two replicates from HT29 and each of the other lines outperformed analysis of five HT29 replicates (Figure 3A). We then considered whether the gRNA numbers could similarly be reduced without sacrificing performance, and evaluated the accuracy of JACKS with 2, 3, 4, or 5 randomly picked gRNAs for each gene in each of the three libraries. While performance decreased with reduction of gRNA numbers, just three randomly selected Avana gRNAs for each gene and two replicates were enough to outperform MaGECK with three replicates and all five gRNAs (Figure 3B). Combined, using two replicates and three gRNAs would reduce the required experiment size 2.5-fold, directly impacting the scale and cost of screens.

**Figure 3:**
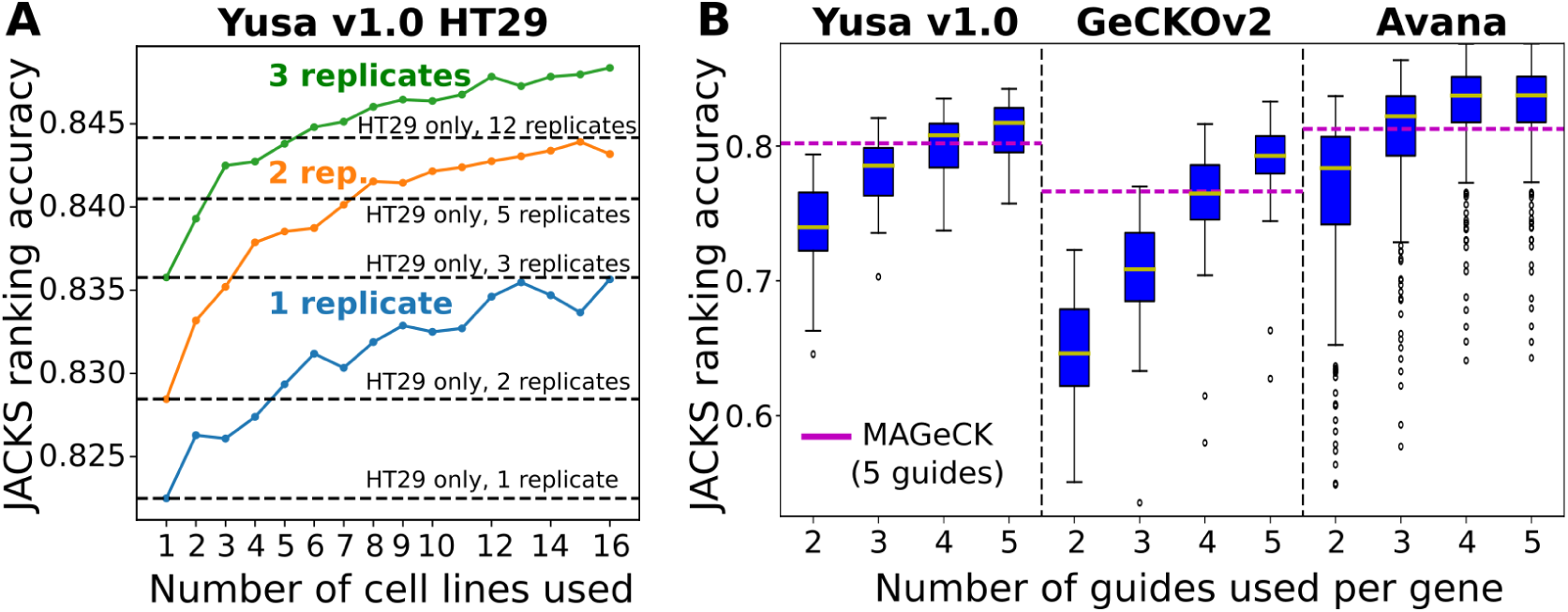
JACKS enables reduced screen size and cost. **A.** Average JACKS ranking accuracy (y-axis) on HT29 cell line for increasing numbers of co-processed cell lines (x-axis), and different number of technical replicates (colors). 200 cell lines samples were randomly generated for each point on the graph, and results averaged. As a reference, the same metric is plotted in increasing numbers of HT29 replicates processed by JACKS without the other cell lines (dashed lines). **B.** JACKS ranking accuracy (y-axis) for increasing numbers of gRNAs (x-axis) from three different libraries (panels) using two replicates per cell line, compared to MAGeCK used on all five gRNAs and all available (2-4 per cell line) replicates (dashed line). Box plot as in Figure 1C.

We presented JACKS, a Bayesian model for joint analysis of screens performed with the same gRNA library. We demonstrated that JACKS improves identification of screen hits compared to existing analysis methods in the vast majority of lines tested across three different data sets and gRNA libraries, and does so without sacrificing context-specific signal. This allows greatly reduced experiment sizes while retaining performance equivalent to MAGeCK, the current analysis standard. Ability to carry out screens with smaller libraries and to efficiently analyze them is especially important for some of the most interesting applications, such as mapping genes in primary cells that can only be obtained in limited numbers, and propagated for a short time.

Copy number variation has been demonstrated to play a role for essential gene inference from CRISPR/Cas9 screens^19, 22, 23, 27^, and is modeled in CERES. While JACKS does not account for this signal distortion, the experiments described here should be independent of copy number effects. JACKS can be used in combination with CRISRPcleanR^22^ or other pre-processing methods that remove these effects.

Recompiling published datasets and re-running a full analysis for each new screen may be prohibitive for many in practice. We have pre-computed gRNA efficacies for these existing libraries, which can be used to process a single screen to achieve equivalent performance to the full JACKS model (Pearson’s *r*^2^ *>* 0.99, Figure S10). This functionality is available via Python, R, and a Shiny web interface.

## Acknowledgements

The authors would like to thank Joshua Dempster for useful discussions about CERES, and Jared Simpson and Oliver Stegle for comments on text. FA was supported by a Royal Commission for the exhibition of 1851 Research Fellowship, LP by Wellcome, and Estonian Research Council (IUT 34-4).

## Author Contributions

FA, LP conceived the method. FA, LP developed the model. FB, FI, KY, and MG contributed the additional Yusa v1.0 screens and provided guidance. FA performed analysis. FA, LP wrote paper with input from all authors.

## Code and data availability

JACKS is available under an MIT license at http://www.github.com/felicityallen/JACKS in Python and R.

## Correspondence

Correspondence and requests for materials should be addressed to FA and LP.

## Methods

### Joint Analysis of CRISPR-Cas9 Knockout Screens (JACKS)

We define the observed log_2_ fold change of the *i*th guide targeting the *g*th gene in the *l*th treatment condition as *y*_*g,i,l*_, where this value is computed as the mean across median-normalized replicate measurements as follows

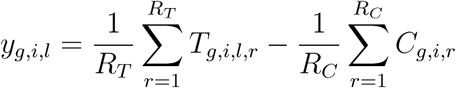

where *T*_*g,i,l,r*_ = log_2_(*t*_*g,i,l,r*_+32)−median(*T*_:,:*,l,r*_) and *C*_*g,i,r*_ = log_2_(*c*_*g,i,r*_+32)−median(*C*_:*,:,r*_) are log-transformations of the raw read counts *t*_*g,i,l,r*_ and *c*_*g,i,r*_ for the *r*th replicates in the treatment and control samples respectively and the median functions operate over all guides across all genes in each respective replicate; *R*_*T*_ and *R*_*C*_ are the number of replicates in those respective samples.

We model *y*_*g,i,l*_ as a Gaussian distribution

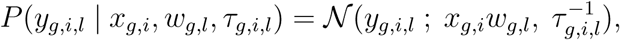

where

- 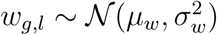 is the condition-dependent gene effect of the *g*th gene in the *l*th treatment condition, where *µ*_*w*_ and 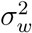 are set to 0 and 1000 respectively for a weak prior that is constant across conditions.
- 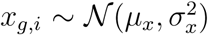 is the condition-independent gRNA efficacy of the *i*th guide targeting the *g*th gene. A stronger prior is specified, with 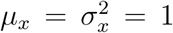 to reflect the prior belief that most gRNAs work moderately well, as well as to prevent over-fitting. To make the model identifiable, the means of the approximate posteriors of *x* are normalized during inference within each gene, such that their median-emphasized average is 1, according to:

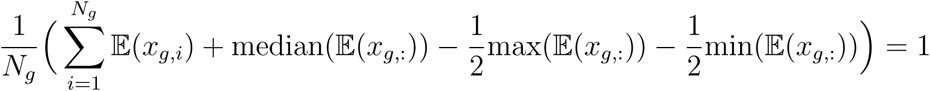

where *N*_*g*_ is the number of gRNAs targeting gene *g*, and *x*_*g*,_: refers to all efficacies for gene *g*. This median-emphasized average is intended to select an appropriate reference point for *w* that accounts for all observations for each gene but up-weights the median and down-weights the extremes.
- *τ*_*g,i,l*_ ~ Γ(*a*_*g,i,l*_, *b*_*g,i,l*_) is the precision of *y*_*g,i,l*_, which uses a non-parametric approach to assign an empirical Bayes prior that accounts for the mean-dependent variability of the log_2_ count values within the replicate measurements of both the treatments and controls. This provides a data-driven and computationally feasible alternative to the parametric approach of modeling counts using a negative binomial distribution, as used in MAGeCK^3^. Given that in general, only 2-4 replicate screens are carried out, direct empirical estimates of these variances are poor. Consequently, we instead compute a smoothed mean-dependent estimate of this empirical variance based on all gRNAs in each condition, and then assign the priors on *τ*_*g,i,l*_ as follows:

1. Compute the mean and variance over replicates for all median-normalized log counts in each treatment and control sample. i.e. the means and variances of *T*_*g,i,l,*_: and *C*_*g,i,*_: where : in the subtexts denote all replicate measurements.
2. Sort these mean-variance pairs by their mean value.
3. Apply a simple moving average filter to the variance values such that each estimated variance becomes a mean of the empirical variances of the 800 gRNAs with closest mean in that cell line (or control), with an additional correction that ensures monotonicity in the relationship (scanning from highest mean, any steps lower are held constant). Denote these estimated variances for each treatment and control measurement as 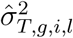 and 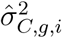 respectively.
4. Assign the prior parameters for *τ*_*g,i,l*_, *a*_*g,i,l*_ = *κ* and 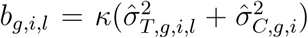, where *κ* determines the strength of the prior (we used *κ* = 0.5), which assigns an expected variance of *y*_*g,i,l*_ as 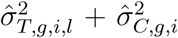, the sum of the estimated treatment and control variances.

Variational inference is used to infer the posterior distributions of *x*, *w* and *τ*. The closed form update equations for the posterior distributions of each variable are:

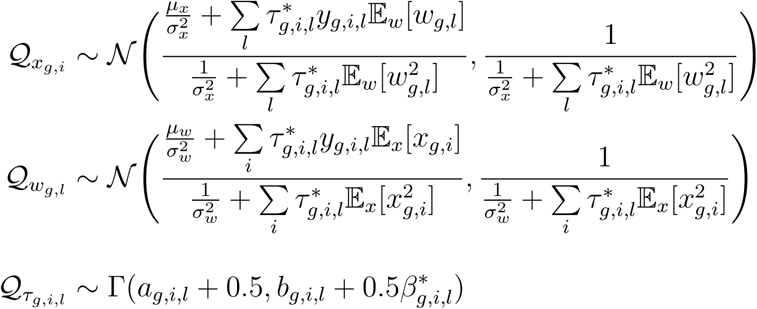

where

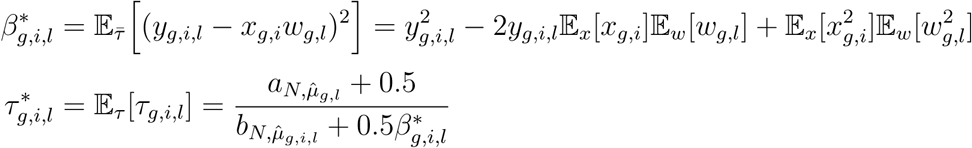

### Classification of Hart essential genes on pooled knock-out screens

The three genome-wide pooled CRSIPR/Cas9 knock-out screen data sets used here were compiled from data in Koike-Yusa *et al.*^28^, Iorio *et al.* ^22^, Aguire *et al.*^19^ and Meyers *et al.*^18^. The compiled sets are listed on our Github repository (see data availability) with complete instructions for recompilation in the README files. We used the BAGEL core essential genes and non-essential gene sets defined by Hart *et al.*^4^, and listed in Github (as above), restricted to those that were targeted by guides within each library. We evaluated performance using the 0.2 partial area under the curve (0.2 pAUC, “ranking accuracy”) and equivalent above the curve (0.2 pAAC, “ranking error”) metrics (Figure 2A, S3). AUCSs are robust measures commonly used to assess the ability of a method to distinguish between two categories, the partial aspect focuses this metric on the more relevant part of the curve where the false positive rate is below 20%. Equivalent results were obtained using other thresholds (0.1 pAUC, full AUC) and performance criteria (recall at fixed false discovery rate, false positive rate at fixed recall, delta AUC (essential vs all genes AUC - non-essential vs all genes AUC); Figure S4-S7, Table S1). All metrics were calculated directly from the receiver operator curve returned by the roc function in scikit-learn^29^ applied to the estimated gene essentiality measures.

### Comparisons with other methods: MAGeCK, BAGEL, CERES, MAGeCK-MLE and MeanFC

Scripts used to run all other methods are available on Github (see Data availability). Input formats for each method were inconsistent, and so data was reformatted for compatibility. MeanFC was computed using a custom script that computed the mean median-normalized log2 fold changes across replicates for each gRNA as done in JACKS (described above) and then assigned each gene a score equal to the mean of this value across all gRNAs targeting that gene. MAGeCK^3^ v0.5.7 test command was used to run MAGeCK and mle command was used to run MAGeCK MLE^27^. BAGEL^4^ v0.91 was slightly modified to take a mean of the control samples (when multiple were available) before computing the fold changes, since BAGEL otherwise expects a single control measurement. CERES v0.0.0.9 was run with *λ*_*g*_ = 0.561 for Avana and *λ*_*g*_ = 0.681 for GeCKOv2 as recommended in Meyers *et al.*^23^. We note that in CERES, *λ*_*g*_ controls the extent to which common gene responses across cell lines are favoured, and so altering this value would alter the results presented here. However, as deciding a correct value for this parameter without over-fitting to the test at hand is non-trivial, we relied on the published recommended values selected for these same datasets, and did not attempt to optimize this further. We believe this is representative of general usage of this program in the absence of alternative guidance, but note that this may be a worthwhile area for future exploration.

### Concordance and reproducibility of gRNA efficacy values

We used the Rule-Set 2 scores from Doench *et al.*^12^ (”Doench-Root Scores”), which provide a sequence-based prediction of gRNA efficacy. Concordance between these scores and JACKS gRNA efficacy estimates was assessed similarly to in Meyers *et al.*^23^, by binning gRNAs by their Doench-Root scores and then looking for increased fractional representation of those gRNAs deemed to have higher *x* value in higher scoring Doench-Root bins. Unlike in Meyers *et al.*^23^, we restricted this analysis to gRNAs targeting Hart essential genes, as we did not expect JACKS (or CERES) to derive any meaningful gRNA efficacy information from genes with no screen activity.

Reproducibility estimates for JACKS gRNA efficacy values were obtained by running JACKS on the Avana batch 0 and batch 1 datasets separately. We computed JACKS gRNA efficacy estimates (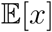), for those gRNAs targeting Hart essential genes^20^, on 100 randomly selected sets of N cell lines from each batch, where N = 1, 3, 5, 8, 10, 15, 20, 25, and 30. We then computed Spearman’s correlation coefficient between the two estimates for each set to obtain a distribution of correlations.

### Additional Yusa v1.0 screens

Previously unpublished screens using the Yusa v1.0 library in HT29, CO205, HuPT4, SW1990 and A375 were carried out using the same screening protocol as in Iorio *et al.*^22^. Raw count data is available on the Github (see data availability).

### Construction of random line data

To generate the five randomly shuffled cell lines for the GeCKOv2 and Avana libraries, we randomly selected three replicates for each line, from existing replicates in other lines. For each of those replicates, we computed the log_2_ median-normalized fold changes as in JACKS method, and then randomly shuffled those fold changes across all guides. The fold changes were then converted back to raw counts, accounting for the control values of their re-assigned gRNAs. The script used to create these lines, and the generated lines are available on the Github (see data availability).

### JACKS with hierarchical prior (HP)

To create a version of JACKS that favours similar gene essentialities across cell lines, we added a hierarchical prior, setting the prior mean *µ*_*w*_ and variance 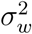 on *w*_*g,l*_ to the current estimated mean of 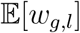 across all cell lines, and three times the current estimated variance of 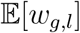, respectively, at each update step in the variational inference. This encourages each *w*_*g,l*_ to be more similar to that in the other lines, with the effect being stronger when there is a more consistent response across lines.

### Compilation of BRAF mutant and wild type cell lines

BRAF mutation status for cell lines in the Avana dataset were obtained from the Cancer Cell Line Encylopedia data portal^30^. Cell lines selected for the melanoma and colon cancer sets were those that were either BRAF WT or BRAF mutant but which did not have amplifications or deletions in the BRAF gene, to avoid issues with copy number differences in comparisons between JACKS and CERES.

### Random sampling to assess impact of the number of cell lines, replicates and gRNAs

To investigate the effect of increasing the number of cell lines co-processed by JACKS (Figure 1C), for a given cell line under test, we bootstrap sampled (with replacement) the requisite number of other cell lines, randomly selecting two replicates from each. We ran JACKS on each set sampled in this manner and recorded the gene scores for the cell line under test. We repeated this 200 times for each test cell line and condition, computing the average ranking accuracy (0.2 partial AUC score) across repetitions for each test cell line. The box plots in Figure 1C show the distribution of these mean scores across cell lines. The same procedure was used to assess the effect of the number of replicates in the Yusa v1.0 HT29 data (Figure 3A), except that the test line was always HT29, and the number of replicates was also altered. This procedure was also used to assess the impact of reducing the number of gRNAs (Figure 3B), except that the full set of cell lines was used in all samples, and the random sampling was instead taken (without replacement) on the available gRNAs for each gene.

## Supplementary Figures

**Figure S1:**
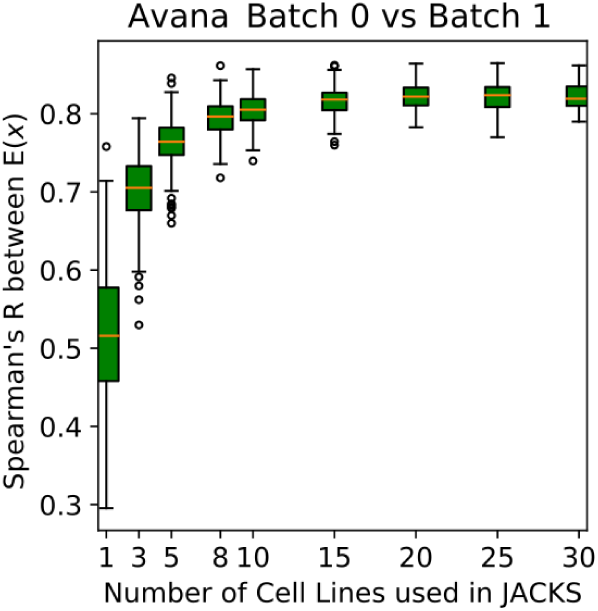
Estimates of gRNA efficacy are reproducible. Spearman’s R (y-axis) for gRNA efficacy estimates from two different batches of Avana library experiments. For increasing number of cell lines used in JACKS (x-axis), 100 random sets of lines were picked for each batch, JACKS was run to infer posterior of gRNA efficacy *x*, and its expected value was used for calculating the correlation.

**Figure S2:**
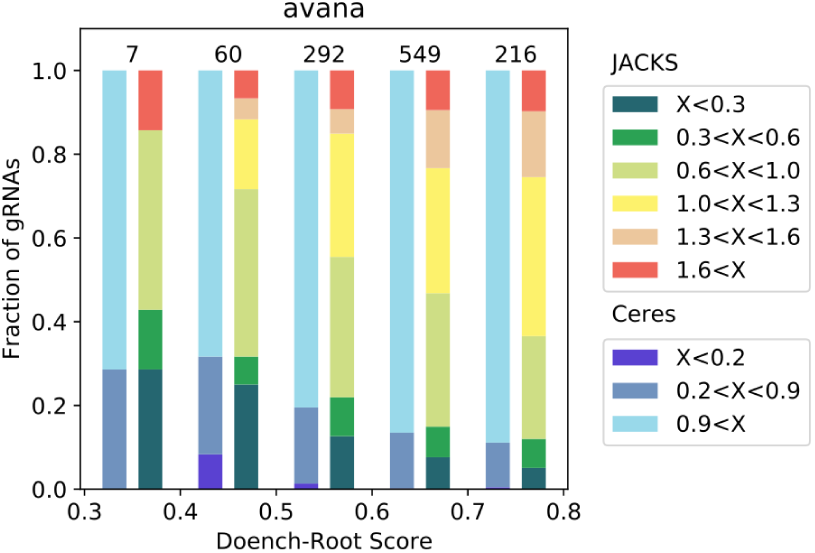
JACKS gRNA efficacy estimate is concordant with CERES estimate and Doench-Root score on the Avana library screens. Fraction of gRNAs (y-axis) in each of the Doench-Root score bins (x-axis) for different strata of CERES and JACKS inferred gRNA efficacies (”x”, colors). Number of gRNAs in each column is marked above the bar.

**Figure S3:**
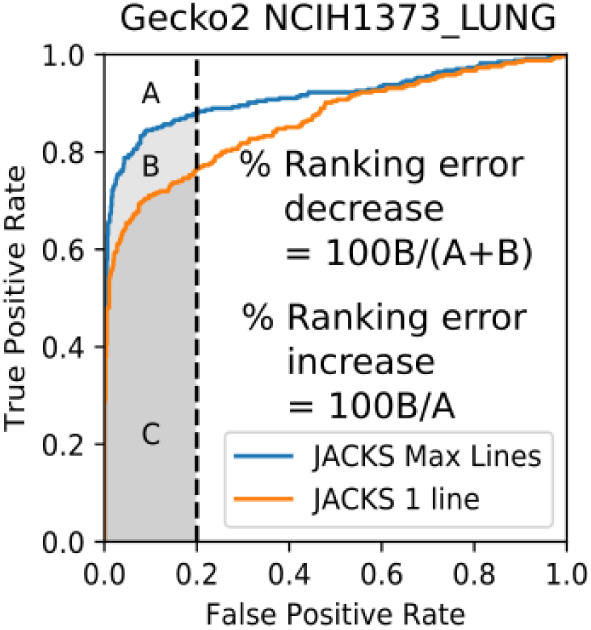
Calculating ranking error and ranking accuracy from receiver operator curve. True positive rate (y-axis) for fixed false positive rate (x-axis) for distinguishing true positive from true negative genes according to Hart *et al.*^4^ using expected gene essentiality from JACKS run on single line (orange) or all 33 GeCKOv2 lines (blue). Dashed line denotes 0.2 false positive rate, ranking accuracy (area under the curve, grey), and ranking error (area above the curve, white) and their change are demonstrated for the NCIH1373 LUNG cell line.

**Figure S4:**
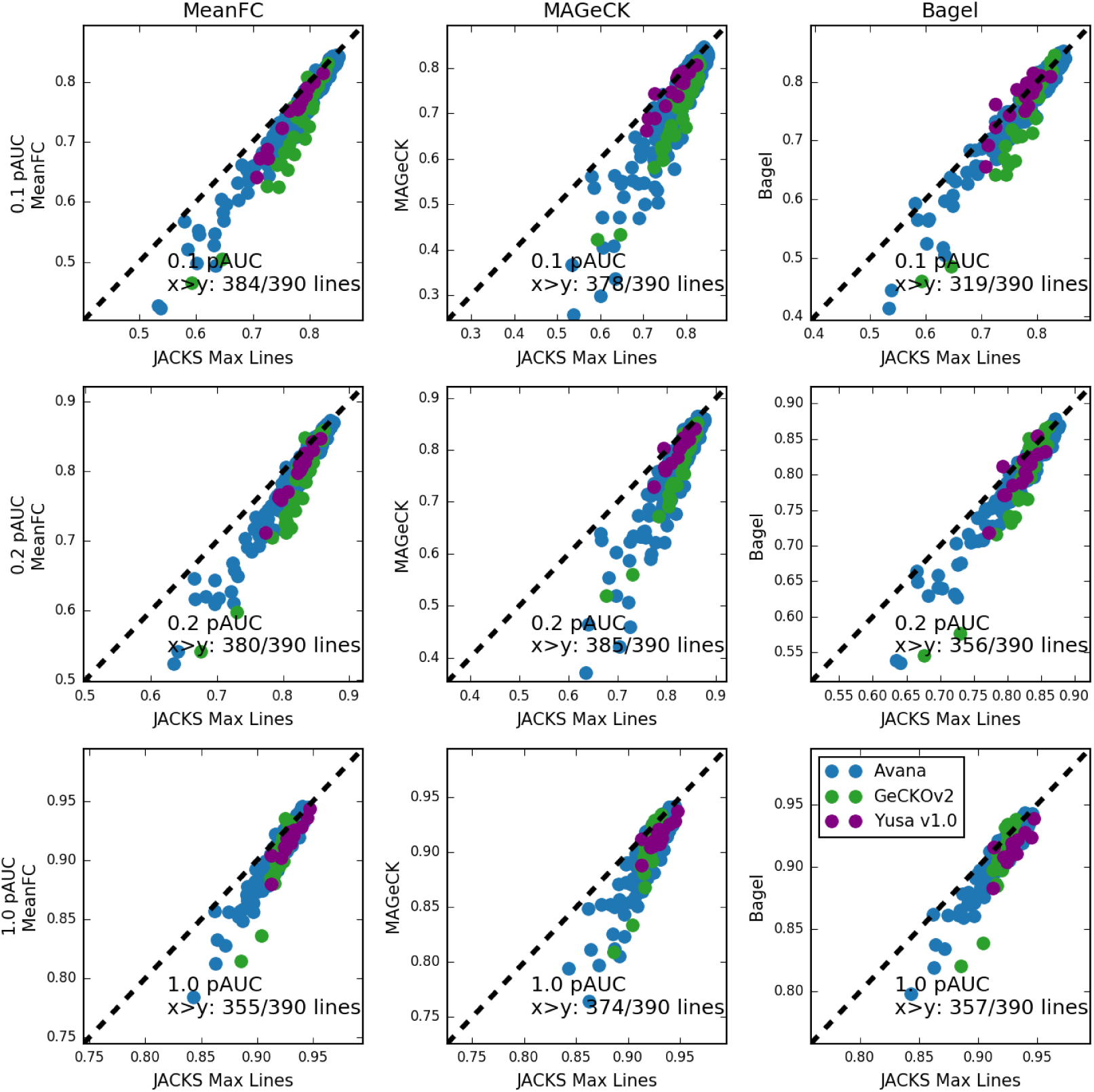
JACKS performs favourably compared to alternatives when measured on area under the curve metrics. JACKS run with all available lines (x-axis) compared to alternative methods (y-axis; first column - mean log2 fold change in gRNA representation; second column - MaGECK p-value; third column - BAGEL essentiality estimate) for cell lines (individual markers) from three different libraries (Avana - blue, GeCKOv2 - green, Yusa v1.0 - purple). First row: partial area under the curve at 0.1 FPR (0.1 pAUC); second row: at 0.2 FPR (0.2 pAUC); third row: at 1.0 FPR (AUC). Number of cell lines for which JACKS has higher pAUC than alternatives is denoted on the plot.

**Figure S5:**
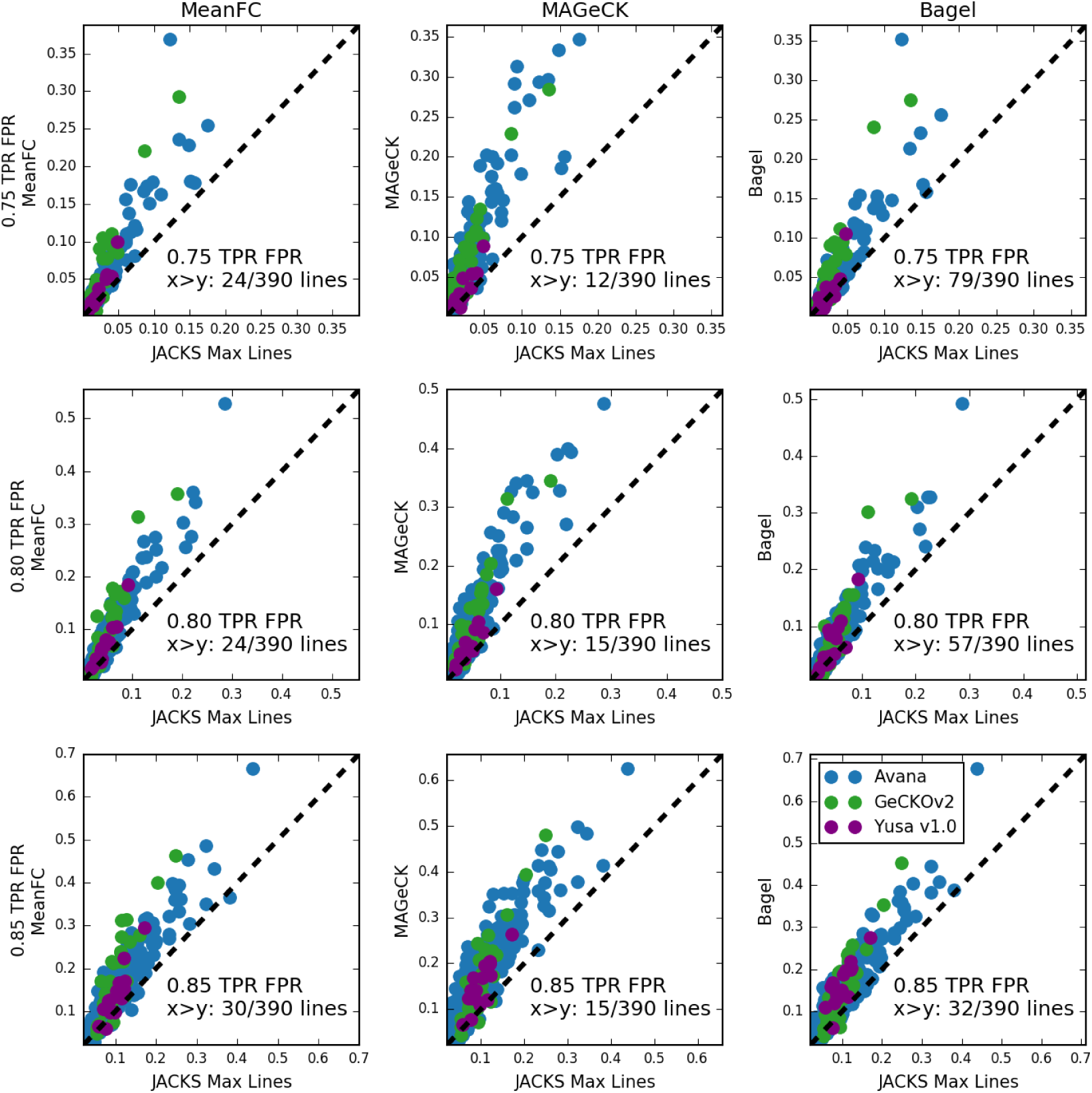
JACKS performs favourably compared to alternatives when measured by false positive rate at fixed recall. Axes, columns, colors as in Figure S4. First row: false positive rate at 0.75 true positive rate (TPR); second row: at 0.80 TPR; third row: at 0.85 TPR. Number of cell lines for which JACKS has higher false positive rate than alternatives is denoted on the plot.

**Figure S6:**
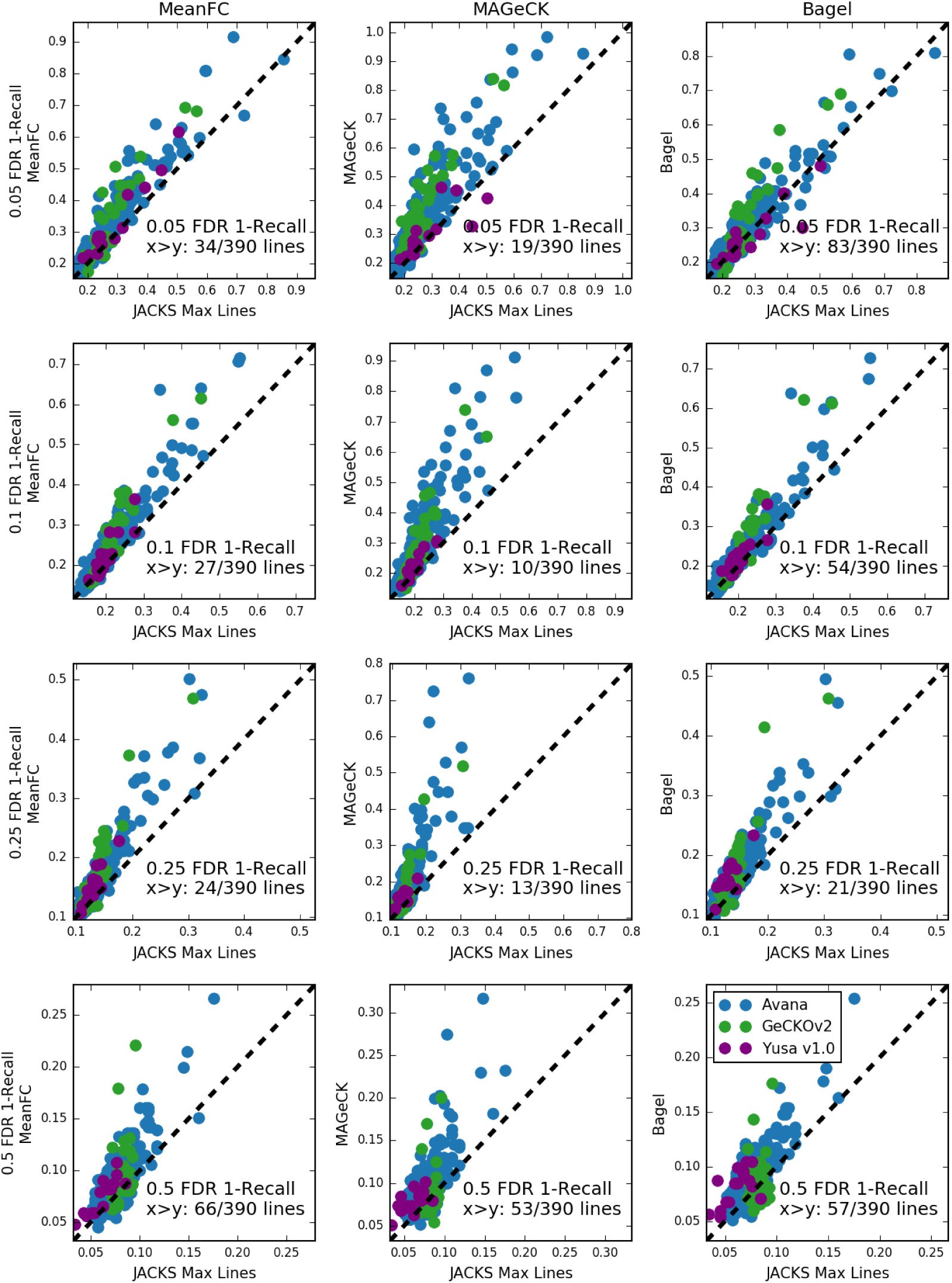
JACKS performs favourably compared to alternatives when measured by false negative rate at fixed false discovery rate. Axes, columns, colors as in Figure S4. First row: false negative rate at 0.05 false discovery rate (FDR); second row: 0.1 FDR; third row: 0.25 FDR; fourth row: 0.5 FDR. Number of cell lines for which JACKS has higher false negative rate than alternatives is denoted on the plot.

**Figure S7:**
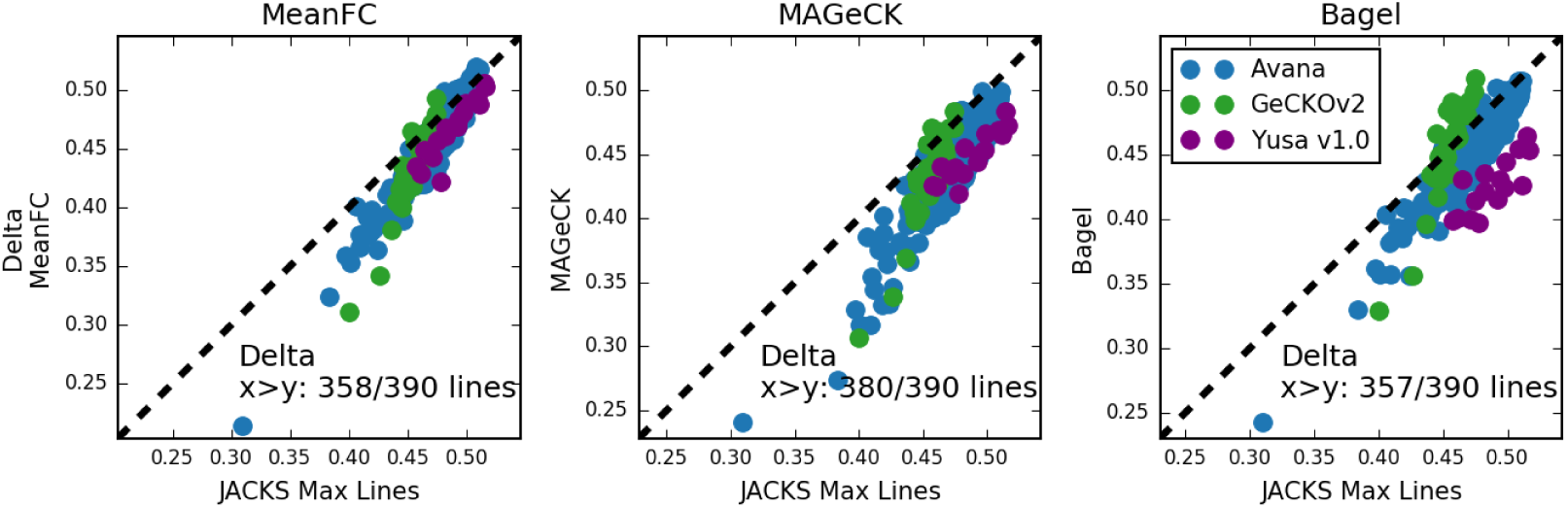
JACKS performs favourably compared to alternatives when measured by delta AUC. Axes, columns, colors as in Figure S4. Number of cell lines for which JACKS has higher improvement in AUC than alternatives is denoted on the plot.

**Figure S8:**
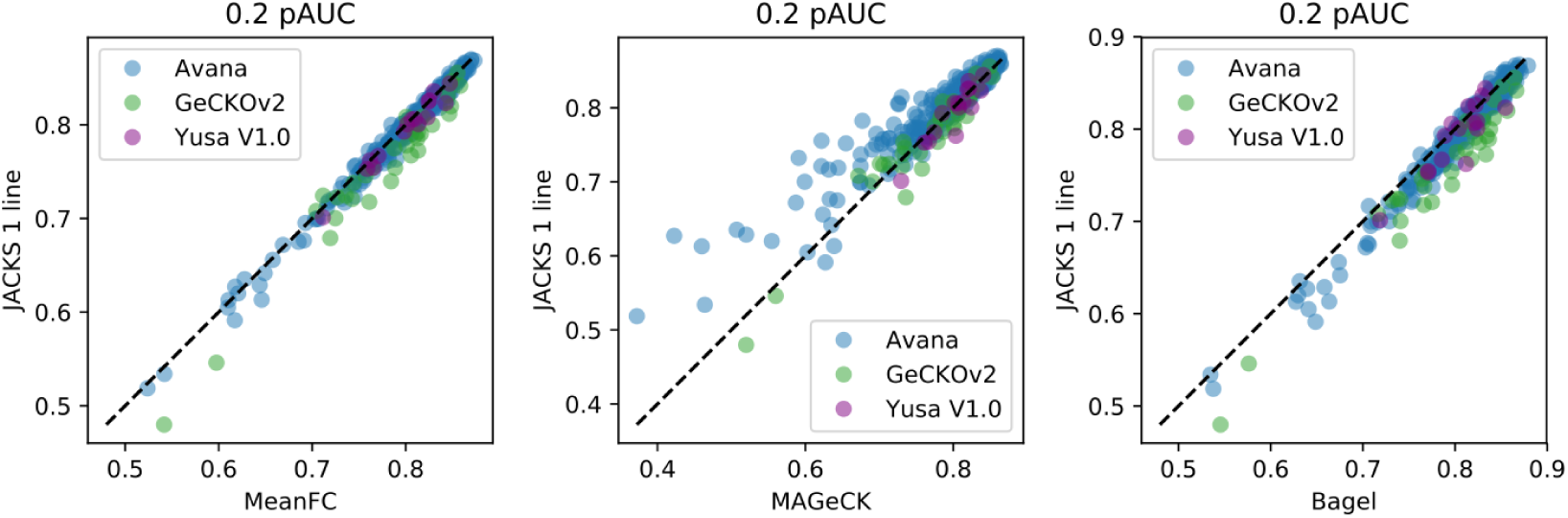
JACKS on only one line at a time performs similarly compared to existing single-line alternatives when measured by parital 0.2 AUC. Axes, columns, colors as in Figure S4.

**Figure S9:**
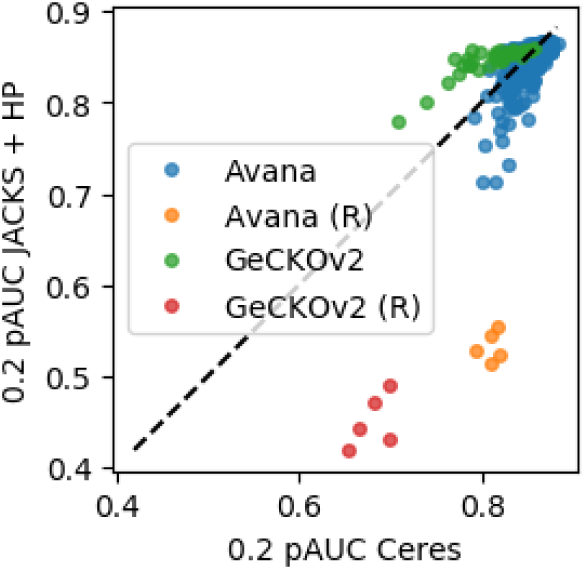
CERES and JACKS(HP) both identify essential genes from random data. Ranking accuracy of CERES (x-axis) compared to JACKS(HP) (y-axis) on Avana (blue) and GeCKOv2 (green) libraries. Each marker corresponds to one cell line, with five randomized experiments (yellow and red markers) included for comparison. Dashed line, *y* = *x*.

**Figure S10:**
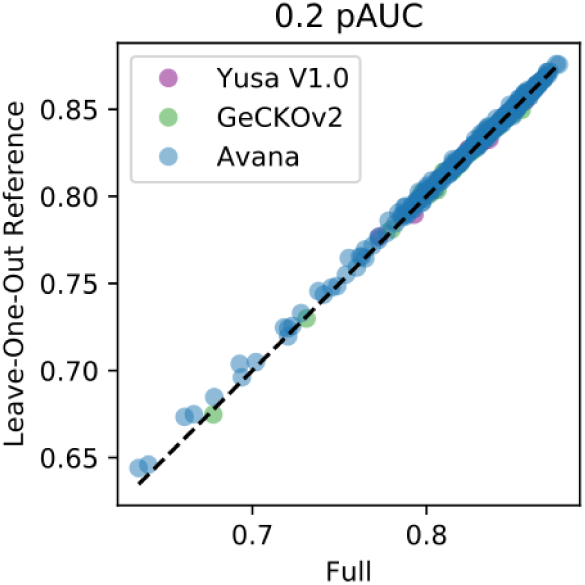
JACKS estimates of gene essentiality can be obtained rapidly using precomputed gRNA efficacies. Recovery accuracy (0.2 pAUC) for full JACKS results (computed on all available data) for a single cell line (x-axis) are nearly identical to those obtained by computing essentialities for a single line at a time, using gRNA efficacies estimated from the rest of the lines (y-axis) for the three considered libraries (colours).

